# Gene expression in cord blood links genetic risk for neurodevelopmental disorders with maternal psychological distress and adverse childhood outcomes

**DOI:** 10.1101/216309

**Authors:** Michael S. Breen, Aliza P. Wingo, Nastassja Koen, Kirsten A. Donald, Mark Nicol, Heather J. Zar, Kerry J. Ressler, Joseph D. Buxbaum, Dan J. Stein

## Abstract

Prenatal exposure to maternal stress and depression has been identified as a risk factor for adverse behavioral and neurodevelopmental outcomes in early childhood. However, the molecular mechanisms through which maternal psychopathology shapes offspring development remain poorly understood. We applied transcriptome-wide screens to 149 umbilical cord blood samples from neonates born to mothers with posttraumatic stress disorder (PTSD; *n*=20), depression (*n*=31) and PTSD with comorbid depression (*n*=13), compared to carefully matched trauma exposed controls (*n*=23) and healthy mothers (*n*=62). Analyses by maternal diagnoses revealed a clear pattern of gene expression signatures distinguishing neonates born to mothers with a history of psychopathology from those without. Co-expression network analysis identified distinct gene expression perturbations across maternal diagnoses, including two depression-related modules implicated in axon-guidance and mRNA stability, as well as two PTSD-related modules implicated in TNF signaling and cellular response to stress. Notably, these disease-related modules were enriched with brain-expressed genes and genetic risk loci for autism spectrum disorder and schizophrenia, which may imply a causal role for impaired developmental outcomes. These molecular alterations preceded changes in clinical measures at twenty-four months, including reductions in cognitive and socio-emotional outcomes in affected infants. Collectively, these findings indicate that prenatal exposure to maternal psychological distress induces neuronal, immunological and behavioral abnormalities in affected offspring and support the search for early biomarkers of exposures to adverse *in utero* environments and the classification of children at risk for impaired development.

## 1. INTRODUCTION

During prenatal development, the fetus is particularly vulnerable to the effects of a broad range of environmental exposures, with consequences that can persist into infancy, adolescence, and adulthood. In particular, maternal psychosocial distress during pregnancy, in the form of exposure to chronic stressors and depression can have profound effects on the developing fetus, and can influence behavioral and physiological outcome measures during infancy and adulthood (DiPietro, 2010; Hart and McMahon, 2006; Kinsella and Monk, 2009; Hollins, 2007). For example, antenatal stress and depression have been linked to behavioral deficits at age five (O’Connor et al., 2013), anxiety disorders at age thirteen (O’Connor et al., 2013; O’Donnell et al., 2014), and have been shown to double the risk of experiencing emotional and behavioral problems for children aged four to seven years (O’Connor, 2002). These exposures have also been associated with reduced infant birth weight (Grote et al., 2010) as well as reductions in head circumference (Koen et al., 2017) and gray matter density in critical cortical regions of the brain (Sandman et al., 2015). While the impact of prenatal maternal distress has been well documented, the potential mechanisms through which maternal psychological variables shape early development have yet to be fully elucidated. An improved understanding of the molecular mechanisms underlying these trans-generational effects is a critical first step in developing strategies for reducing the disease burden on both mother and child.

Increasing evidence suggests that stress-vulnerability established *in utero*, as a result of excessive exposure to maternal hormones *(i.e.* glucocorticoids) and inflammatory cytokines, may ultimately lead to increased risk of psychopathology in the offspring (Lupien et al., 2009; Knuesel et al., 2014; Estes and McAllister, 2016). For example, several studies have leveraged umbilical cord blood (UCB), which represents neonatal blood, to examine the epigenetic-based determinants underlying this hypothesis. Differences in UCB DNA methylation have been reported, including loci encoding for glucocorticoid receptors (Oberlander et al., 2008), serotonin transporters (Devlin et al., 2010), immune system-related genes (Nemoda et al., 2015) and neuronal development-related genes (Lillycrop et al., 2015; Sparrow et al., 2016; Starnawska et al., 2017); all of which have been associated with prenatal exposure to maternal anxiety and depression *(see review*, Hodyl et al., 2016). In parallel, experimental animal models have begun to demonstrate the biological plausibility of a causal effect, showing that maternal immune activation, as a consequence of maternal viral infection or neuropsychiatric disorders, is sufficient to impart lifelong neuropathology and altered behaviors in affected offspring (Knuesel et al., 2014; Rovonsky et al., 2015; Estes and McAllister, 2016).

These studies also highlight several fundamental questions. First, while it is clear that epigenetic mechanisms regulate gene expression and underlie the trans-generational effects of maternal psychological stressors (Tsankova et al., 2010), studies have not explored the hypothesis that maternal psychological distress is associated with UCB transcriptional signatures implicating stress-response, immune signaling and neuronal development-related genes. Second, there is a paucity of human studies investigating the trans-generational effects of maternal post-traumatic stress disorder (PTSD) on UCB, and ultimately how UCB gene signatures compare between neonates born to mothers with PTSD and depression. Third, if the evidence supports altered gene expression patterns as a consequence of prenatal exposure to maternal PTSD and depression, it is then important to elucidate whether these changes in gene expression also precede clinically recognizable deficits in physiological and developmental measures in affected offspring. Fourth, maternal psychological distress, traumatic events and its sequelae are highly prevalent low- and middle income (LMIC) settings such as South Africa, and there are very few data emerging from these regions; there is therefore a critical need to examine whether prenatal exposure to maternal psychological distress predicts alterations in UCB and/or physiological and behavioral deficits in young children, particularly in the LMIC setting.

The goal of the current study was to begin to fill these gaps in knowledge by leveraging our longitudinal prospective mother-infant cohort, the Drakenstein Child Health Study based in South Africa. We analyzed transcriptome-wide gene expression profiles of 149 UCB samples from neonates born to mothers with prenatal PTSD (n=20), depression (n=31) and PTSD with comorbid depression (PTSD/Dep; n=13), compared to neonates born to carefully matched trauma exposed controls without meeting PTSD criteria (TE; n=23) and healthy mothers (n=62). We also evaluated physiological and developmental measures in these infants at birth, six months and twenty-four months. A multistep analytic approach was used that specifically sought to: 1) identify dysregulated genes, molecular pathways and discrete groups of co-regulated gene modules in UCB associated with prenatal maternal psychopathologies; and 2) to determine the impact of perinatal PTSD and depression on early childhood development outcomes.

## 2. MATERIALS AND METHODS

### 2.1. Participants and study design

Our study at the Cape Town site is a nested sub-study within a longitudinal birth cohort project, the Drakenstein Child Health Study, which aims to investigate the impact of early life exposures on development of childhood health and disease (Zar et al., 2014; Stein et al., 2015). Participants were pregnant women recruited from community clinics (Mbekweni, TC Newman), and Paarl Hospital of the Drakenstein sub-district, South Africa. This site was selected for its stable population, low socioeconomic status, and free public health system, all of which is representative of many peri-urban regions in South Africa and other low- and middle-income countries. Pregnant women in the study were enrolled at 20-28 weeks gestation followed prospectively and longitudinally to 5 years of age. Maternal diagnoses and data included in the current study were collected antenatally at 28-32 weeks gestation (*i.e.* third trimester) and postnatally at 16 weeks, 6, 18 and 24 months. A combination of standard self-report measures on trauma exposure and psychological symptoms and clinician-administered interviews were given at each time point. Likewise, their infants were followed for 24 months, including comprehensive clinical assessments at 24 months using the Bayley Scales of Infant and Toddler Development, 3^rd^ edition (Bayley-III) (Bode et al., 2014; Weiss et al., 2010; Bayley, 2005).

### 2.2. Prenatal maternal diagnoses

Prenatal maternal trauma exposure was defined as either *i)* directly experiencing a traumatic life event, *ii)* learning that a close family member or close friend experienced a traumatic life event, or *iii)* losing a family member or close friend to illness or other causes. Maternal PTSD during the prenatal period was assessed with the modified PTSD Symptom Scale (mPSS), a psychometrically validated, 18-item self-report scale with excellent internal consistency, high test-retest reliability, and concurrent validity, as well as mirroring the DSM-IV criteria for PTSD (Foa et al., 1993). Thus, women with a history of trauma exposure and who met DSM-IV criteria for PTSD based on the mPSS during the prenatal period were categorized as prenatal PTSD cases. Likewise, women with a history of trauma exposure but did not meet criteria for PTSD during the prenatal period based on the mPSS were considered trauma-exposed (TE) controls. Prenatal maternal depression was assessed with the Beck Depression Inventory II (BDI-II) (Beck and Steer, 1984; Dutton et al., 2004) around 28 to 32 weeks of gestation. A BDI-II cutoff score of 14 yielded a sensitivity of 88% and specificity of 84% when compared to a diagnosis of major depressive disorder (MDD) made by PRIME-MD Mood Module, a focused interview for MDD using DSM-IV criteria. Hence, participants with prenatal BDI score ≥15 were grouped as prenatal maternal depression. Women with a history of trauma exposure who met criteria for PTSD based on the mPSS, and had a BDI score ≥15 during the prenatal period were considered having comorbid PTSD and depression (PTSD/Dep). Participants with no history of trauma exposure or depression were categorized as healthy controls.

Prenatal maternal tobacco use was measured with the tobacco subscale of the Alcohol, Smoking and Substance Involvement Screening Test (ASSIST), which was validated in a multi-site international study, as well as in Cape Town, South Africa (Humniuk et al., 2008; van der Westhuizen et al., 2016). Women with a score of 0 on the tobacco subscale of ASSIST were categorized as non-smokers while those with a score above 0 were considered smokers. Prenatal maternal alcohol use was measured using the alcohol subscale of the ASSIST (*i.e.* women were classified as ‘alcohol-exposed’ if they scored ≥11) or if they gave a positive history on the alcohol exposure questionnaire (AEQ) of alcohol use in any of the three trimesters of pregnancy at levels consistent with WHO moderate-severe alcohol use (*i.e.* either drinking two or more times a week or two or more drinks per occasion). Alcohol exposure for two women with no available ASSIST or AEQ data was assessed using the Alcohol Use Disorder module of the Mini-International neuropsychiatric interview (MINI).

### 2.3. Infant clinical measures

The Bayley-III is a comprehensive assessment of infant development in children aged 0-3 years (Bode et al., 2014; Weiss et al., 2010; Bayley, 2005). As a gold standard measure of infant development globally, the Bayley-III has been widely used in LMIC settings. The tool consists of five scales of infant development: *i)* cognitive; *ii)* communication (receptive and expressive); *iii)* motor (gross and fine); *iv)* socio-emotional (*e.g.* self-regulation, communicating needs, engaging others, using emotional gestures/signals to solve problems); v) adaptive behavior, and assesses development using direct observation of the infant as well as caregiver report. The adaptive behavior scale assesses daily functional skills of the child which are necessary for increasing independence and to meet environmental demands. Areas assessed are communication (speech, language, listening, nonverbal communication); community use (interest in the activities outside the home and recognition of different facilities); health and safety (showing caution and keeping out of physical danger), leisure (playing, following rules, and engaging in recreation at home); self-care (eating, toileting, bathing); self-direction (selfcontrol, following directions, making choices); functional pre-academics (letter recognition, counting, drawing simple objects); home living (helping adults with household tasks and taking care of personal possessions); social (getting along with other people - using manners, assisting others, recognizing emotions); and motor (locomotion and manipulation of the environment). Each scale is scored according to the Bayley-III manual using specialized software (*Bayley-III Scoring Assistant Update Version 2.0.2 with Bayley-III PDA conduit).* Scaled scores are calculated using age-specific reference norms. Each subscale has a possible score range of 1-19. In this cohort, the Bayley-III is administered at twenty-four months of age.

### 2.4. Cord blood collection, RNA isolation and microarray hybridization

Umbilical cord blood (UCB) was collected by trained staff after offspring delivery but before delivery of the placenta. The cord was clamped and cut, after which the clamp was released and cord blood drained by gravity into a kidney collection dish. Thereafter, cord blood was collected using a syringe for processing and storage at −80°C in PAXgene RNA tubes. Staff were specifically trained to ensure only cord blood drained into the collection dish and that to their best ability, no other/external blood was collected. Only samples of offspring whose mothers had provided informed consent for the collection, storage, and future analyses of RNA were eligible for inclusion.

Subsequently, RNA was extracted from PAXgene RNA tubes. All 168 samples had Bioanalyzer RNA Integrity Number (RIN) ≥7. Complementary DNA was derived from RNA and then hybridized, and raw probe intensities were generated on Illumina HumanHT-12 v4 BeadChip arrays. These 168 samples were randomized with respect to sex, maternal diagnosis, maternal alcohol and tobacco use, and mode of delivery (*i.e.* natural delivery, elective C-section, or emergent C-section) on 14 chips to reduce the chance of batch effects.

### 2.5. Gene expression data processing, quality control and differential expression

All statistical analyses were conducted in the statistical package R. All gene expression data were processed, normalized and quality treated. Raw data were background corrected, quantile normalized and log_2_ transformed using the *neqc* function of limma (Ritchie et al., 2015), a robust method for Illumina BeadChips. Non-specific filtering retained probes expressed in at least 50 arrays with a detection P-value <0.05. When multiple microarray probes mapped to the same HGNC symbol, we used a median reduction technique and used the resulting collapsed expression profile for each gene for further analysis. Resulting normalized data were inspected for outlying samples using unsupervised hierarchical clustering of samples and principal component analysis to identify potential outliers outside two standard deviations from these grand averages. Based on these metrics, 19 samples were labeled as outliers and 149 samples passed into the subsequent analyses. Linear mixed models from the R package varianceParition (Hoffman and Schadt, 2016), were used to characterize and identify biological and/or technical drivers that may affect the observed gene expression. This approach quantifies the main sources of variation in each expression dataset attributable to differences in biological factors (*e.g.* maternal diagnoses, age, sex ethnicity), environmental factors (*e.g.* alcohol and tobacco use, clinical medication, mode of delivery) and technical factors (*e.g.* RIN, batch).

The frequencies of circulating immune cells were estimated for each quality controlled sample using Cibersort cell type de-convolution (Newman et al., 2015) (https://cibersort.stanford.edu/). Cibersort relies on known cell subset specific marker genes and applies linear support vector regression, a machine learning approach highly robust compared to other methods with respect to noise, unknown mixture content and closely related cell types. As input, we used the LM22 signature matrix to distinguish 9 main immune cell types: B cells, cytotoxic T cells (CD8+), helper- and regulatory T cells (CD4+), dendritic cells, macrophages (M0-M2), mast cells, natural killer (NK) cells (CD56+), monocytes (CD14+) and neutrophils. The LM22 matrix can be further divided into 13 less frequent immune cell subsets, which we pooled and defined as ‘other’. The resulting estimates were tested for normality using Kolmogorov-Smirnov test and a Dunnett’s multiple comparison of means test with post hoc Tukey correction was used assess differences between neonates exposed to prenatal maternal psychopathologies (*e.g.* PTSD, depression, TE) with neonates born to healthy mothers. A family-wise error rate P < 0.05 was considered a significant difference when performing multiple comparisons of prenatal exposure groups with controls.

Differential gene expression (DGE) analysis was applied to detect relationships between distinct maternal diagnoses and UCB gene expression levels using the limma package (Ritchie et al., 2015). The covariates maternal tobacco use, maternal alcohol use, RIN, batch, biological sex, gestational age, mode of delivery and ethnicity were included in all models to adjust for their potential confounding influence on UCB gene expression between main group effects, and the significance threshold was a nominal P-value <0.05. This P-value threshold was used to yield a sufficient number of genes to include within functional annotation and gene network analyses, described below. Power and sample size was estimated using the R package ssize.fdr (Orr and Liu, 2015).

### 2.6. Weighted gene co-expression network analysis

Weighted gene co-expression network analysis (Langfelder and Horvath, 2008) (WGCNA) was used to build a signed co-expression network using a total of 10,705 genes. To increase confidence and power to detect biologically meaningful modules, a consensus network was built using all 149 samples, as described previously (Breen et al., 2015). The absolute values of Pearson correlation coefficients were calculated for all possible gene pairs and resulting values were transformed with an exponential weight (β=7) so that the final matrix followed an approximate scale-free topology (R^2^>0.80). The WGCNA dynamic tree-cut algorithm was used to detect network modules (minimum module size=30; cut tree height=0.99; deep-split=3, merge module height=0.25). Once network modules were identified, modules were assessed for significant associations to distinct maternal diagnoses, as well as other biological and technical factors. In order to determine which modules, and corresponding biological processes, were most associated with maternal diagnostic states, we ran singular value decomposition of each module’s expression matrix and used the resulting module eigengene (ME), equivalent to the first principal component, to represent the overall expression profiles for each module. Gene significance (*GS*) was also computed as the -log_10_ of the *P*-value generated for each gene based on linear regression analyses (described above in differential gene expression), and is a measure of the strength of differential gene expression between each comparison being tested. Module significance (*MS*) was calculated as the average *GS* within each module. Differential analysis of MEs was performed using a ANOVA testing with *post hoc* Tukey correction. Finally, for each gene in a module, intramodular membership (*kME*) was defined as the correlation between gene expression values and *ME* expression. Genes with high *kME* inside co-expression modules are labeled as hub genes.

### 2.7. Enrichment analyses

All DGE signatures (*P*<0.05) and module genes with high connectivity (kME>0.7) were assessed for functional enrichment using three complementary methods. First, DGE and module enrichment categories were obtained using REACTOME pathways from the publically available Enrichr software (Chen et al., 2013; Kuleshov et al., 2016) (*http://amp.pharm.mssm.edu/Enrichr/*) and *p*-values were computed using a hypergeometric test assuming a binominal distribution and independence for probability of any gene belonging to any gene set. Second, we downloaded the highly expressed, cell specific gene expression database compiled by Shoemaker *et al.* 2012, as well as brain cell type markers from Zeisel *et al.* 2015, to assess modules for over-representation of cell type specific markers. Third, neurodevelopmental disorder genetic risk loci were curated from human whole-exome and genome-wide association studies of autism spectrum disorder (ASD) (Sanders et al., 2015), developmental delay (Gene2Phenotype, OMIM, 2017), intellectual disability (Parikshak et al., 2013), schizophrenia (Fromer et al., 2014) and epilepsy (EuroEPINOMICS-RES Consortium, 2014). We also curated lists of gene coexpression modules found to be dysregulated in the cerebral cortex of autistic cases (Voineagu et al., 2011) and differentially expressed genes in umbilical cord blood from neonates born to mothers with socio-economic disadvantage (Miller et al., 2017). Overrepresentation analysis was determined using a one-sided Fisher’s exact test with Benjamini-Hochberg multiple test correction.

### 2.8. Computational code and data availability

All computational code for quality control of gene expression profiles, differential gene expression analysis and weighted gene co-expression network analysis are available through request to the corresponding author and freely accessible from the following site: *https://github.com/BreenMS/Umbilical-cord-blood-transcriptome-analysis*. Raw expression data are freely available at the Gene expression Omnibus (GEO) under accession number GSEXXXXX (*accession will be released following manuscript acceptance*).

## 3. RESULTS

This exploratory study applied transcriptome-wide screens to 149 UCB samples from neonates born to mothers with posttraumatic stress disorder (PTSD; *n*=20), depression (*n*=31) and PTSD with comorbid depression (PTSD/Dep; *n*=13), compared to carefully matched trauma exposed controls (TE; *n*=23) and healthy mothers unexposed to trauma without depression (*n*=62) (**Table 1**). A multi-step analytic approach was used that specifically sought to 1) identify individual genes and 2) functional molecular pathways (*i.e.* co-expression modules) associated with distinct maternal psychopathologies, and 3) determine the impact of prenatal exposure to maternal psychopathology on early childhood development outcome measures.

### 3.1. Neonate group-wise comparisons by maternal diagnosis

Following standardized data pre-processing procedures, the proportions of circulating immune cells were estimated for all individuals since complete cell counts with leukocyte differentials were not available. Comparative analyses of the estimated immune cell type proportions showed no significant differences between neonatal groups (**Supplementary Figure 1**), suggesting that immune cell type frequencies would not confound downstream analyses. Subsequently, we applied differential gene expression (DGE) analysis to identify UCB expression signatures associated with prenatal exposure to maternal depression, PTSD, PTSD/Dep and TE, using neonates born to healthy mothers as a baseline reference for all comparisons (**Supplementary Table 1**). The overall relatedness of these four lists of covariate-adjusted DGE signatures were examined by constructing a matrix of log_2_ fold-changes from these comparisons and a Jaccard coefficient to create a direction of effect gene-based phylogeny. Gene expression profiles clustered accordingly to prenatal exposures, with similar patterns of expression observed for neonates exposed to TE and depression, followed by depression and PTSD/Dep, and finally PTSD/Dep and PTSD (**Fig. 1A**). Scatterplots and pairwise overlaps of the DGE lists were used to further quantify this result and identified significant overlaps between: prenatal PTSD and PTSD/Dep groups (∩=35, *P*=0.009); prenatal depression and PTSD groups (∩=65, *P*=0.8); and prenatal PTSD and TE groups (∩=67, *P*=0.002) (**Fig. 1B-D**). Interestingly, DGE signatures between prenatal PTSD and TE groups displayed a strong negative correlation when compared to healthy controls (*r*=-0.50, *P*=2.5E-99) (**Fig. 1D**), indicating that gene expression profiles respond in an opposite direction for prenatal TE compared to PTSD, suggestive of PTSD resiliency signatures. These differences pressed us to directly compare prenatal PTSD and TE groups, for which we found a number of under-expressed genes in PTSD (over-expressed in TE) significantly related to (*P*<0.05) corticotropin-releasing hormone (*CYP11A1, FOSB, RAF1*) and circadian rhythm (*NOS2, LEP, CSNK1E*) and over-expressed genes in PTSD (under-expressed in TE) related to cytoskeleton remodeling (*RSU1, LIMS1*) and interferon signaling (*OAS1, STAT2, XAF1*) (**Supplementary Table 2**). Over-expressed genes related to prenatal PTSD/Dep were similarly involved in interferon signaling (*OASI, STATI, STAT2, XAF1, IFIT3, IFIT2*) and under-expressed genes were involved in gluconeogenesis (*TPI1, GOT2, ALDOA, GAPDH*). In comparison, over-expressed genes related to prenatal depression were implicated in translation and mRNA splicing and under-expressed genes were involved in *NCAM1* interactions. All functional enrichment analyses based on DGE lists can be found in **Supplementary Table 2**. Overall, all of the identified DGE signatures were mainly associated with a prenatal exposure to distinct maternal psychopathologies and not with any other confounding factors, including age, sex, mode of delivery, nicotine, or estimated cell type frequencies (**Supplementary Figure 2**).

### 3.2. Differential module expression and prenatal maternal depression

We applied weighted gene co-expression network analysis (WGCNA) to identify discrete groups of co-regulated genes (modules), and identified 14 modules (**Supplementary Figure 3**). Subsequently, all modules were tested for overrepresentation of differentially expressed genes related to prenatal maternal depression. Two modules implicated in axon guidance (M2) and mRNA stability (M13) were strongly enriched for prenatal depression DGE signatures, as reflected by elevated module significance (MS) values (**Fig. 2A**). To determine whether the overall expression of these modules was significantly associated to prenatal depression, we calculated differences in module expression using module eigenene (ME) values (*See Materials and Methods for complete description of ME*). Consistent with results using *MS*, ME expression for M2 was under-expressed and ME expression for M13 was over-expressed in the depression group compared to healthy controls (*P*=0.03; *P*=0.05, respectively) (**Fig. 2B,C**). Subsequently, ME values were correlated to all clinical parameters found in **Table 1**, to determine module-trait relationships. Both modules M2 and M13 were significantly associated with prenatal depression (**Supplementary Table 3**) and were not correlated to any other measured clinical variable, suggesting that differential gene expression in these modules was not confounded by technical or environmental variables (*e.g.* mode of birth, medication, tobacco *etc*.). Cell type enrichment analysis revealed that the decreased axon guidance-related module M2 was strongly enriched for neuronal and astrocyte cell type signatures, whereas the increased mRNA stability-related module M13 was enriched for innate and adaptive immune cell types (**Fig. 3A,B**). Hub genes for M2 included *GARNL4, LDB2, MGAT4B* and *ISL2*, and hub genes for M13 included *ALDOB, IGFBP7, IFIT2*, and *LOC727913* (**Supplementary Figure 4**). Interestingly, the genes with the highest intramodular connectivity (kME>0.7) in module M2 displayed a negative association with prenatal depression (Pearson’s *r*=-0.38, P=1.5E-20) that increased steadily across PTSD/Dep, PTSD and TE diagnoses (**Supplementary Figure 5A**).

### 3.3. Differential module expression and prenatal maternal PTSD

Additionally, two modules were significantly over-represented with prenatal PTSD DGE signatures relative to TE, including module M3 involved in tumor necrosis factor (TNF) signaling as well as module M14 involved in cellular response to stress (**Fig. 2D**). ME expression for M3 was significantly under-expressed and ME expression for module M14 was significantly over-expressed in the PTSD group relative to TE (*P*=0.005; *P*=0.03, respectively) (**Fig. 2E,F**). Similarly, ME values were significantly associated with prenatal PTSD and not with other confounding factors (**Supplementary Table 3**). Cell type enrichment revealed that the decreased module M3 was strongly enriched for CD4+ and CD8+ T cell and microglial cell type markers, whereas the increased module M14 was enriched for CD71+ early erythroid cell markers, and to a lesser extent, oligodendrocyte cell type markers (**Fig. 3A,B**). Hub genes for module M3 include *GPX8, SIGLEC6, TMEM173* and *BCYRN1* and hub genes for module M14 include *WWC3, EP300, TXNRD1* and *COPA* (**Supplementary Figure 6**). Overall, no modules were associated to prenatal PTSD/Dep, perhaps due to its moderate sample size (*n*=13).

### 3.4. Mapping neurodevelopmental disorder genetic risk loci onto disease modules

We next sought to determine whether any of the disease-related modules also contained a significant over-representation of neurodevelopmental disorder genetic risk variants identified from previous large-scale whole-exome and genome-wide association studies, which may imply a further causal role for altered neurodevelopmental outcomes in offspring. Surprisingly, we found that three out of the four disease-related modules contained a significant enrichment for neurodevelopmental risk loci (**Fig. 3C**). Prenatal depression module M13, involved in mRNA stability, contained a significant overrepresentation of developmental delay genetic risk loci (∩=71, *P*=0.02). In parallel, prenatal PTSD module M3, involved in TNF signaling, was enriched for schizophrenia genetic risk loci (∩=37, *P*=0.01), and cellular response to stress module M14 was also enriched for autism spectrum disorder (ASD) genetic risk loci (∩=9, *P*=0.0009), including the highly penetrant *CHD8* locus. Module M14 also displayed a significant overlap with a previously identified gene co-expression module found to be significantly dysregulated in the cerebral cortex of ASD individuals (∩=25, *P*=0.02) as well as with previously identified DGE signatures in UCB from neonates born to mothers with socio-economic disadvantage (∩=120, *P*=<0.001) (**Fig. 3C, Supplementary Figure 7D**). These significant enrichments corroborate the fact that these modules are independent of technical or environmental effects and represent likely primary disease effects. Overlapping genes and risk variants are listed in **Supplementary Table 4** and their intramodular network positions are described in **Supplementary Figure 6**.

### 3.5. Clinical follow-up measures in affected infants

Importantly, we assessed whether developmental outcome measures in neonates exposed to prenatal maternal psychological stressors differed from those born to healthy mothers. A standardized developmental assessment (Bayley-III) at twenty-four months and physiological measures were collected at birth, six months and twenty-four months for a subset of infants for which gene expression was measured (PTSD, *n*=14; depression, *n*=27; PTSD/Dep, *n*=8; TE, *n*=14; healthy controls, *n*=48). While no differences were observed for physiological measures between groups (**Supplementary Table 5**), we identified significantly reduced cognitive performance at twenty-four months for infants exposed to prenatal PTSD, depression and TE compared to healthy controls (*P*=0.04, *P*<0.001, *P*=0.04, respectively) (**Fig. 4A, Supplementary Table 6**). In addition, we also observed significant reductions in social-emotional outcomes for infants exposed to prenatal TE and PTSD/Dep compared to healthy controls (*P*=0.04, *P*=0.04, respectively) (**Fig. 4B**) as well as reduced performance on adaptive behaviour subscales, including community use scores, at twenty-four months for infants exposed to prenatal PTSD/Dep compared to healthy controls (*P*=0.02) (**Fig. 4C**).

## 4. DISCUSSION

In this investigation we utilize UCB gene expression data collected from the South African Drakenstein Child Health Study to identify genes and molecular pathways associated with prenatal exposure to distinct maternal psychopathologies. We observed gene expression patterns distinguishing between neonatal groups, as well as distinct expression profiles in neonates echoing PTSD resiliency signatures when mothers were resilient to PTSD after trauma exposure. We identified two discrete groups of co-regulated genes (modules) that relate to prenatal depression and two modules that relate to prenatal PTSD, leading us to focus on these modules. Two of these modules were strongly enriched for brain-expressed genes, including prenatal depression module M2 and prenatal PTSD module M14. Using data from a previously published transcriptome, whole-exome and genome-wide association studies, the overall genes represented in three-disease modules were enriched with genetic risk loci for developmental delay (M13), schizophrenia (M3) and ASD (M14), which further support the primary role of these modules in impaired developmental outcomes. Importantly, these gene expression relationships (at both gene and modular levels) were independent of demographic, environmental or technical factors. Finally, the dysregulation of these molecular pathways preceded reductions in cognitive and socio-emotional performance as well as in adaptive behaviour in a subset of affected infants. We discuss these points in turn below.

A central finding was the identification of gene modules associated to distinct psychopathologies found in our maternal sample. Two modules were associated with prenatal exposure to maternal depression, including module M2 implicated in axon guidance and enriched for neuronal cell types and module M13 involved in mRNA stability and enriched for genetic risk loci implicated in developmental delay. In parallel, two modules were associated with prenatal exposure to maternal PTSD, including module M3 implicated in TNF signaling and enriched for schizophrenia risk loci, and module M14 implicated in cellular response to stress and enriched for ASD genetic risk loci, as well as immune and inflammatory genes found differentially expressed in UCB of neonates born to mothers with socio-economic disadvantage (Miller et al., 2017) as well as in the cerebral cortex of ASD cases (Voineagu et al., 2011). The enrichments with brain cell types and neurodevelopmental disorder genetic risk loci corroborates the fact that these modules are independent of technical effects and likely represent primary disease effects. Previous studies finding enrichment of disease related genetic variants in their co-expression modules of interest, implicated that these modules represent causal effects (Voineagu et al., 2011; de Jong et al., 2012; Greenawalt et al., 2011). Both ASD and schizophrenia have been associated with increased microglial and immune activity in postmortem tissue (Voineagu et al., 2011; Morgan et al., 2010; Radewicz et al., 2000) as well as elevated levels of inflammatory cytokines in peripheral blood (Tylee et al., 2016; Ashwood et al., 2008; Michel et al., 2012). There is also evidence suggesting that symptom severity for both disorders correlate with the immune system dysfunction (Michel et al., 2012; Song et al., 2009), and since immune abnormalities endure into adulthood and anti-inflammatory agents appear to be a beneficial intervention (Song et al., 2009; Rossignol et al., 2009), it is possible that these immune changes may actively contribute to disease symptoms. Importantly, pre-clinical animal studies have provided further evidence that prenatal maternal immune activation alters neurodevelopment and leads to behavioral changes that are relevant for ASD and schizophrenia (Golan et al., 2005; Meyer et al., 2006; Boksa 2010; Kneeland and Fatemi, 2013). Collectively, our data align with these studies to suggest that neurodevelopmental immune insults and genetic background interact and may ultimately increase risk for either disorder. Our approach also highlights a methodological strength, in that network analyses can be used to reconstruct molecular phenotypes for the identification of the genetic association signal derived from pathways, rather than small effects from individual genes.

Another interesting novel finding from our DGE analysis was the identification of an inverse correlation of gene expression profiles between prenatal TE and PTSD (**Fig. 1D**), indicating that when compared to neonates born to healthy mothers, neonates with prenatal TE respond in an opposite direction than those with prenatal PTSD. When directly comparing prenatal TE and PTSD groups, over-expressed genes in neonates with prenatal PTSD related to the regulation of cytoskeleton remodeling and interferon signaling and under-expressed genes related to corticotrophin-releasing hormone and circadian rhythm (*P*<0.05). Interestingly, among the significant dysregulated genes with inverse correlations were *CREB1, DMC1* and *MSH3* (over-expressed in TE) and *CAMK2A, NUP160* and *SNW1* (over-expressed in PTSD). These genes, especially *CREB1* and *CAMK2A*, are critical components for the phosphorylation of CREB, a transcription factor that regulates several down-stream targets involved in PTSD pathophysiology, including brain derived neurotropic factor (Galter and Unsicker, 2000a, 2000b), circadian clock genes (Yehuda et al., 2005) and numerous neuropeptides (Reul and Holsboer, 2002) (*e.g.* corticotrophin-releasing hormone); all of which have been implicated in epigenetic studies investigating early life adversity in cord blood, placental tissue and maternal venous blood (Nemoda et al., 2015; Sparrow et al., 2016; Kertes et al., 2017). Moreover, when compared to neonates born to healthy mothers, several prenatal PTSD/Dep-related genes also related to increased interferon signaling, indicating a degree of molecular specificity for heightened interferon signaling in UCB following prenatal exposures to PTSD and PTSD/Dep, relative TE and healthy controls. Comparably, a recent blood transcriptome mega-analysis also found a high degree of transcriptional dysregulation across biological sex and modes of trauma in PTSD, which converged on interferon signaling pathways (Breen et al., 2017). Candidate gene studies have similarly found considerable evidence for increased interferon gamma underlying the onset and emergence of PTSD symptoms (Passos et al., 2015), which when taken together support the heighted interferon signature in the current study. Moreover, when compared to healthy controls, over-expressed genes in neonates with prenatal TE predominantly related to non-canonical NF-κβ signaling (*LTBR, TNDSF12, PSME2, PSMB1, PSMB8, BIRC2, BIRC3, SKP1*) and several over-expressed prenatal PTSD genes were implicated in TNF-α signaling (*NPAS, IKBG, TANK, BCL2L1*), a proinflammatory cytokine that ultimately activates canonical NF-κβ signaling. While the exact cause and effect relationships stimulating interferon genes in PTSD and PTSD/Dep remain unresolved, our results imply that trauma exposure triggers classical stress related pathways that are detectable across UCB and adult peripheral blood, and depending on the severity of the psychological burden, may activate interferon signaling (as in PTSD onset) or potentially trigger alternative non-canonical inflammatory pathways through suppressed TNF-α expression (as in PTSD resiliency).

These results raise several fundamental questions. First, what is the relevance of UCB gene expression for assessing brain mechanisms in offspring? UCB represents neonatal blood and has been a critical tissue for examining alterations in neurotransmitter and inflammatory markers, and potential dysregulation of the *in utero* environment. Interestingly, a study investigating the short- and long-term effects of maternal depression on DNA methylation patterns found a significant number of overlapping differentially methylated loci across both neonate UCB and adult hippocampi, many of which were involved in immune system function (Nemoda et al., 2015). Previous research into the correspondence of gene expression across blood and brain compartments reveals weak to moderately strong correlations between tissues (Tylee et al., 2013). However, if the biomarker is in association with an adverse *in utero* event or a poor neurodevelopmental outcome, its expression may not, or necessarily need to, resemble the expression of the same analyte in the brain. Second, what is the clinical utility of UCB gene expression for identifying newborns exposed to adverse *in utero* events? Our diagnostic panel was able to distinguish healthy neonates from those exposed to prenatal maternal stressors with 73% on withheld test data, indicating that RNA-based biomarkers may be clinically useful but need to be validated once larger sample sizes become available. Third, does prenatal exposure to maternal depression and PTSD increase subsequent risk for ASD and schizophrenia in offspring? Although somewhat inconsistent, there are data to suggest that exposures to severe stressors early in pregnancy or even immediately prior to conception increase the risk of schizophrenia primarily in male offspring (Khashan et al., 2008), while bereavement stress during the third trimester is associated with an increased risk for ASD (Class et al., 2014). As mentioned above, mounting evidence indicates that both ASD and schizophrenia share genetic risk factors related to immune function, which suggest that maternal psychological distress, perhaps interacting with alterations in immune function, may contribute towards increased disease risk. In this light, the altered neuro-immune processes as a result of maternal psychological distress reported in this study may possibly potentiate the effects of neurodevelopmental risk genes-an idea that is supported by preclinical studies demonstrating marked interactions between maternal immune activation and neurodevelopmental risk (Estes and McAllister, 2016; Morgan et al., 2010; Radewicz et al., 2000; Drexhage et al., 2010). Fourth, are the most informative disease genes central (‘hub’) network genes or do they display low intramodular connectivity? We find that many of the genes that have been linked to previous genetic risk variants or transcriptomic investigations reside on the periphery of disease-related gene co-expression modules (**Supplementary Table 4**), similar to previous reports (Gaiteri and Sibille, 2011). The low kME for the majority of these genes could imply that modulating candidate disease genes for therapeutic intervention is likely to have a limited effect due to their non-central network location, perhaps accounting for the low efficacy of current treatments. Finally, how do our findings inform disease mechanisms through which maternal psychopathology shape offspring development? Our findings run parallel with pre-clinical studies that suggest maternal distress can alter axon guidance, inflammatory and cellular response to stress pathways in UCB, which aggravate postnatal development, including reduced cognitive scores and adaptive behaviors at two years of age.

Several limitations of our study deserve emphasis. First, while heighted cortisol and inflammation are hallmark signatures of PTSD and depression, we did not have antenatal maternal molecular data, which could directly support the maternal immune activation hypothesis. Second, as delayed cognitive performance is a key clinical feature of ASD and has also been shown to predict SCZ, we are unable to provide any direct causal links between maternal diagnoses and the eventual onset of either of these disorders since we did not collect any structured diagnostic information for ASD or SCZ. Third, the observed changes in developmental performance in the cognitive domain may potentially be related to other factors, including maternal withdrawal or less effective parenting skills. Additional caution in over-interpreting these findings is warranted, as UCB contains a mixture of different cell types, for which we provide *in silico* cell type frequency estimates, and some of the reported findings were detected using a nominal *P*-value cut-off (*P*<0.05). Power and sample size calculations indicate that in order to achieve 80% power to detect DGE signatures at a FDR of 0.10, sample sizes in each maternal group must be between 51 to 67 cases and controls (**Supplementary Figure 8**). Since our samples comprise slightly less than 51 to 67 cases and controls, the power of our dataset may be slightly lower than the estimated power. To overcome any limitations related to sample size and to increase our ability to determine signal from noise in a biologically meaningful way, we focused on the identification of groups of coordinately expressed genes (modules). We and others have used WGNCA to identify variable dependency, and have taken advantage of this information to simply mainstream analysis, in terms of heavily reducing the number of multiple comparisons (*e.g.* from >10K genes to tens of modules) and facilitating the interpretation of large data (*e.g.* by focusing on modules with likely biological origins). To add additional safeguards to our analysis and interpretation, we focused on modules meeting all of the following criteria: 1) those significantly associated with maternal depression or PTSD; 2) those with significant functional enrichment terms; 3) those related to the underlying cellular architecture; 4) those which were also significantly enriched with previously defined neurodevelopmental genetic risk loci. In this fashion, we found substantial supporting genetic and genomic evidence for our disease-related genes and gene modules. Future large-scale initiatives combining genetic and functional genomic data from both mother and infant, with detailed antenatal clinical report information, may provide a more detailed mechanism of action for a specific course of impaired neurodevelopment in affected offspring.

In summary, our exploratory study provides a framework for investigating previously unknown molecular aspects of maternal psychological distress on UCB gene expression, which links altered neuro-immune UCB gene expression profiles to maternal depression, PTSD and PTSD resilient phenotypes on the developing fetus. It is hoped that this study will lead to future work confirming the importance of the observed differences in axon guidance, TNF signaling and cellular stress response, and their involvement in shaping developmental outcomes in affected infants. Moreover, the enrichment of brain expressed genes and genetic risk variants with the disease-related gene modules suggests that maternal psychological distress may potentiate the effects of neurodevelopmental risk genes via altered expression of regulatory genes in these networks. Future studies involving suitable model systems could aim to validate these causal hypotheses. Ideally, these findings will contribute towards advanced early biomarkers, which could help target impaired developmental outcomes in affected offspring through early behavioral or pharmacological intervention.

## Author contributions

DS, AW, KD and HZ provided principal oversight for tissue collection, gene expression generation, clinical and developmental data collection and experimental design. AW and NK provided oversight for curating clinical data. MSB formulated the research question, analyzed and interpreted the data and wrote the manuscript. All authors contributed to editing the manuscript and approved the final version.

## Acknowledgements

We thank the Drakenstein Child Health Study staff, the clinical and administrative staff of the Western Cape Health Department at Paarl Hospital and at the clinics and the families and children who participate in this study. We also thank Dr. Anna Tocheva for critical reading of our manuscript.

---

**Figure 1.**
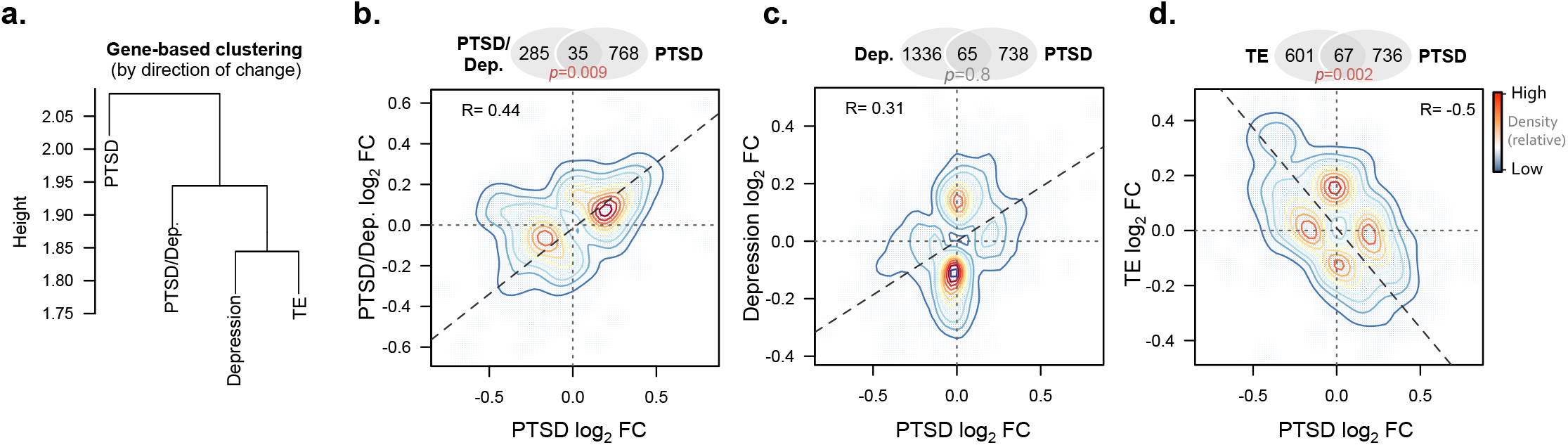
Concordance between differential gene expression results, **(a)** Clustering of gene-based log_2_ fold-changes (direction of change) using four lists of DGE signatures from neonates with distinct prenatal exposures. Neonates born to healthy mothers were used a baseline reference for all comparisons. Correspondence between neonates born to mothers with PTSD relative to neonates born to mothers with **(b)** PTSD/Dep, **(c)** depression and **(d)** trauma exposure (TE) without PTSD. Overlaps and correlations are displayed for all significantly dysregulated genes (*P*<0.05) and contours indicate the density of the overlap. Abbreviations: PTSD, posttraumatic stress disorder; TE, trauma exposure without PTSD; PTSD/Dep, PTSD with comorbid depression.

**Figure 2.**
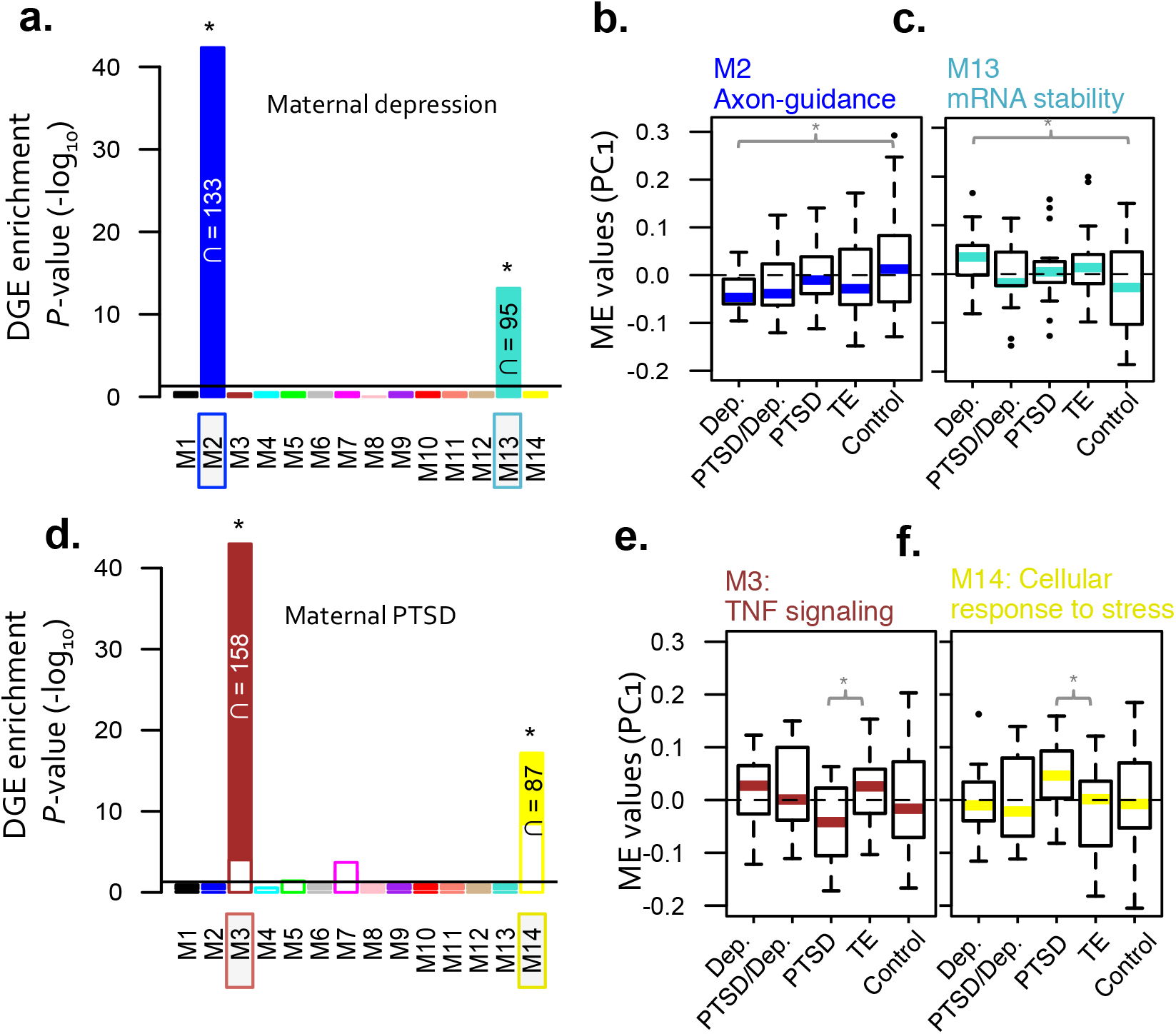
Weighted gene co-expression network analysis of 149 cord blood transcriptome profiles (*n*^genes^=10,705). Module significance (MS) was calculated by testing for over-representation of disease-related differential gene expression signatures (*P*<0.01) within each module and identified **(a)** two depression-related modules and **(d)** two PTSD-related modules. Modules denoted with an asterisk (*) have ME values significantly correlated to conditional states (i.e. maternal depression or PTSD). Representative modules with high MS were investigated for module expression differences using a ANOVA testing and a P-value < 0.05 was considered significant. Boxplots are displayed for each main group. Significant differences in ME expression were observed in maternal depression-rlatd modules **(b)** M2 and **(c)** M13 and maternal PTSD-related modules **(e)** M3 and **(f)** M14.

**Figure 3.**
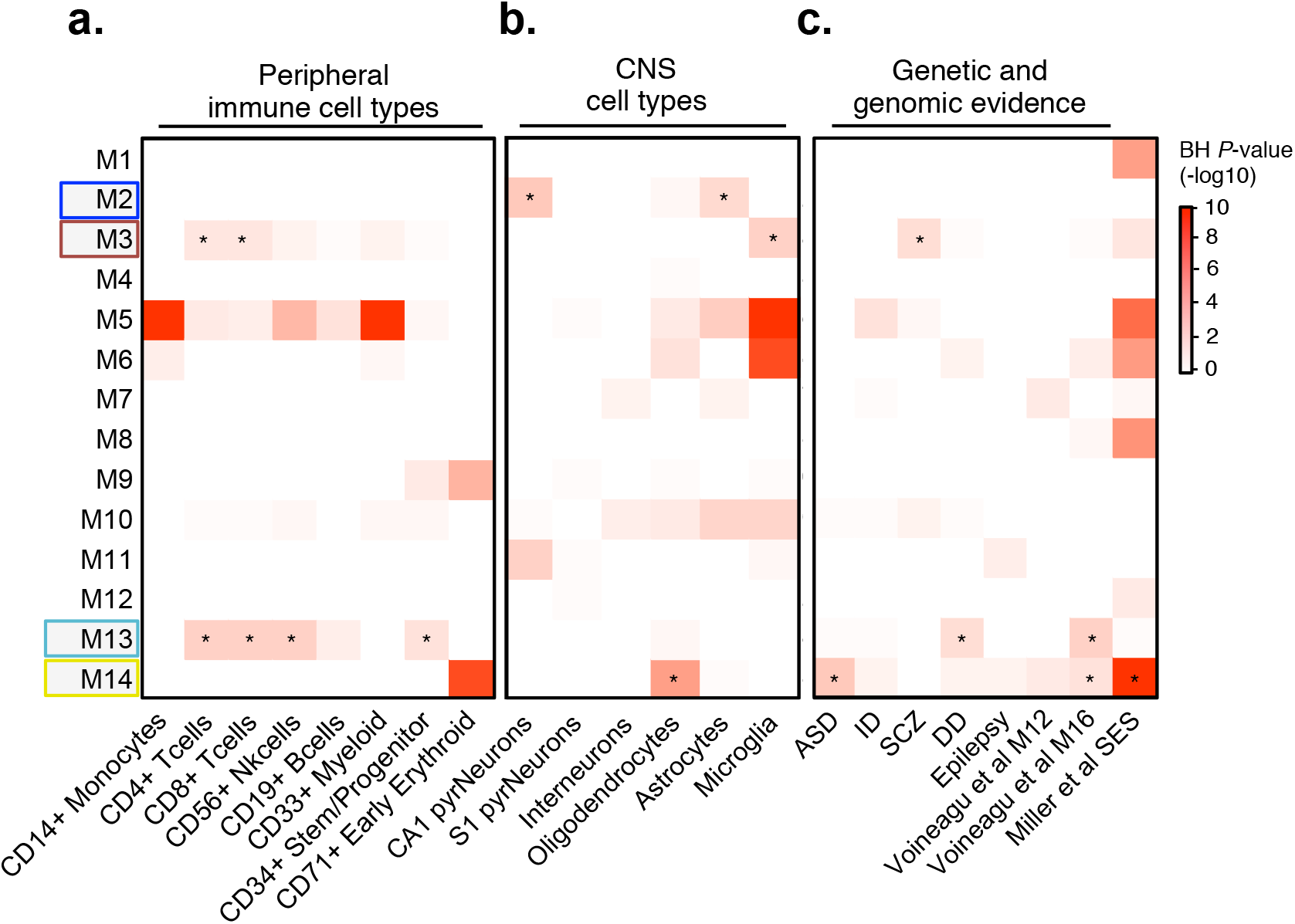
Cell type and genetic enrichment analysis. Modules were tested for enrichment of **(a)** peripheral immune cell types, **(b)** central nervous systems (CNS) cell types, and **(c)** previously identified neurodevelopmental genetic risk loci and genomic risk evidence. Voinague et al., 2011 refers to an altered gene module in the cerebral cortex of ASD cases (M16 from the study) and Miller et al., 2017 refers to UCB gene expression profiles associated with maternal socioeconomic disadvantage. Overrepresentation analysis of these gene sets within network modules was analyzed using a one-sided Fishers exact test to assess the statistical significance. All *P*-values, from all gene sets and modules, were adjusted for multiple testing using Benjamini-Hochberg procedure. We required an adjusted *P*-value < 0.05 (*) to claim that a gene set is enriched within a user-defined list of genes. Modules enriched for DGE signatures are boxed along the y-axis. Abbreviations: ASD, autism spectrum disorder; ID, intellectual disability; SCZ, schizophrenia; DD, developmental delay.

**Figure 4.**
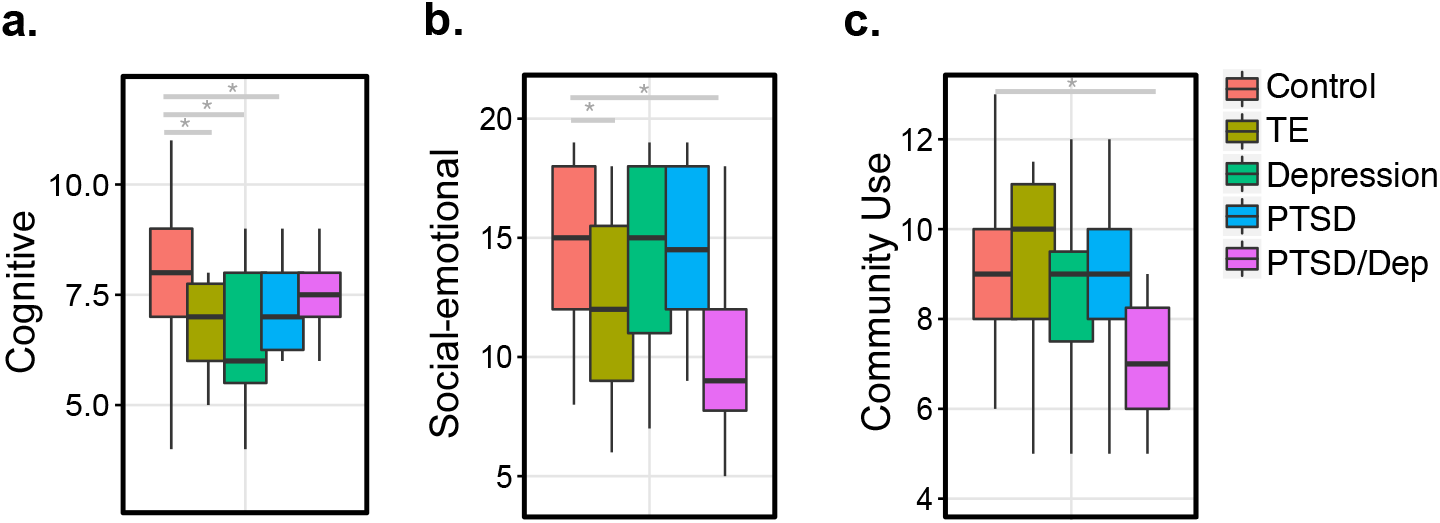
Clinical data were collected for a subset of neonates (*n*= 115), which underwent gene expression analysis. Differences in Bayley-III **(a)** cognitive scores, **(b)** socio-emotional scores and **(c)** adaptive behavior sub-scores (that is, community use) at twenty-four months. Dunnet's multiple comparisons of means statistical testing compared neonatal groups with prenatal exposures to adverse events (*i.e*. PTSD, depression, PTSD/Dep and TE) to healthy mothers, while covarying for nicotine and alcohol. Adjusted *p*-values are reported within brackets using a single-step procedure.

**Table 1.**
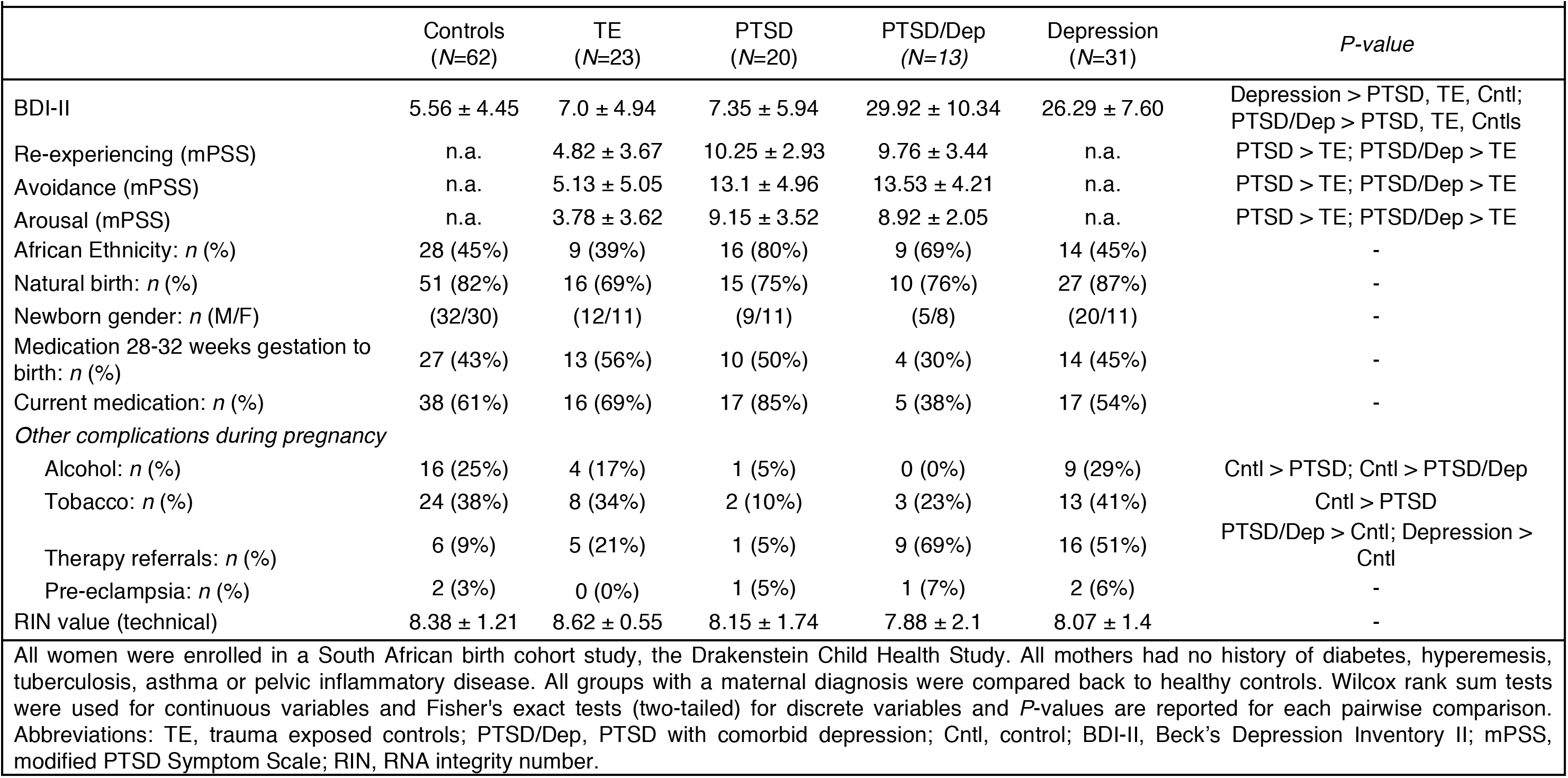
Recorded clinical measures for all participants (*N*=149).

